# The Tube-Sealing Test: A simple assay that reveals offspring-centered defensive behavior in postpartum mice

**DOI:** 10.64898/2025.12.24.696449

**Authors:** Noriko Horii-Hayashi, Shoma Miki, Hanano Takashima, Koshiro Akaki, Koichi Inoue

## Abstract

Perinatal mental health is a major public health concern, and postpartum women are particularly vulnerable to conditions such as postpartum obsessive–compulsive disorder and postpartum depression. Many postpartum symptoms involve heightened concern for infant safety, suggesting the presence of offspring-centered defensive processes that are distinct from general anxiety focused on self-safety. One offspring-centered defensive behavior described in wild rats is entrance-sealing, in which lactating females plug the entrance of their burrow to limit access by potential intruders. Although laboratory mice rarely exhibit sealing spontaneously, sealing-like behavior can be experimentally induced by chemogenetic activation of hypothalamic perifornical Urocortin-3 neurons, indicating that mice retain a latent motor pattern for this response. However, no method has existed for reliably quantifying sealing behavior under controlled laboratory conditions. To address this gap, we developed the Tube-Sealing Test (TST), an assay that measures bedding packed into tube-like openings attached to the home cage and enables repeated, non-invasive quantification of sealing behavior. Using this approach, we found that tube-sealing occurred rarely in males, intermittently in virgin females, and with markedly greater intensity in postpartum females. Although virgin and postpartum females showed a similar likelihood of performing sealing, postpartum females inserted substantially more bedding than either group. In postpartum females, exposure to a male intruder increased sealing specifically at the tube through which the intruder was introduced, indicating that sealing intensity is modulated by direct threat experience. Together, these findings establish the TST as a simple and reproducible method for quantifying sealing behavior and identify tube-sealing as a measurable component of offspring-centered defensive behavior.

## 1. Introduction

Perinatal mental health is a major public health concern, and postpartum women are particularly vulnerable to mental disorders such as postpartum obsessive–compulsive disorder and postpartum depression (Hillerer et al., 2011; Russell et al., 2013; Perani and Slattery, 2014; Meltzer-Brody et al., 2018; Ferra et al., 2024) . Many postpartum symptoms involve heightened concern for infant safety, suggesting the presence of offspring-centered defensive processes, which are conceptually distinct from general anxiety focused on self-safety.

In rodent research, postpartum anxiety is typically examined by exposing mothers to stressors—such as restraint, pup separation, or chronic stress—and then assessing behavior in assays like the open field test and elevated plus maze test (Neumann, 2003; Perani and Slattery, 2014; Yang et al., 2015). However, these assays primarily measure self-focused anxiety, which reflects the animal’s own safety, and therefore do not capture defensive responses specifically oriented toward offspring protection. As a result, the mechanisms underlying offspring-centered anxiety and defense remain poorly understood.

One offspring-centered defensive strategy documented in wild rodents is *entrance-sealing*, in which lactating females block the entrance of their burrow to limit access by potential intruders (Calhoun, 1963) (Fig. 1A). Although laboratory mice do not build burrows under standard housing conditions, sealing-like behavior can be induced experimentally through chemogenetic activation of Urocortin-3 (UCN3) neurons in the perifornical area of the anterior hypothalamus (PeFA) (Horii-Hayashi et al., 2023) (Fig. 1B), indicating that mice possess a latent motor pattern for sealing. Spontaneous sealing in laboratory mice, however, is rare and highly variable, although it can occasionally be observed in postpartum females (Fig. 1C), making it difficult to study under controlled conditions.

**Figure 1.**
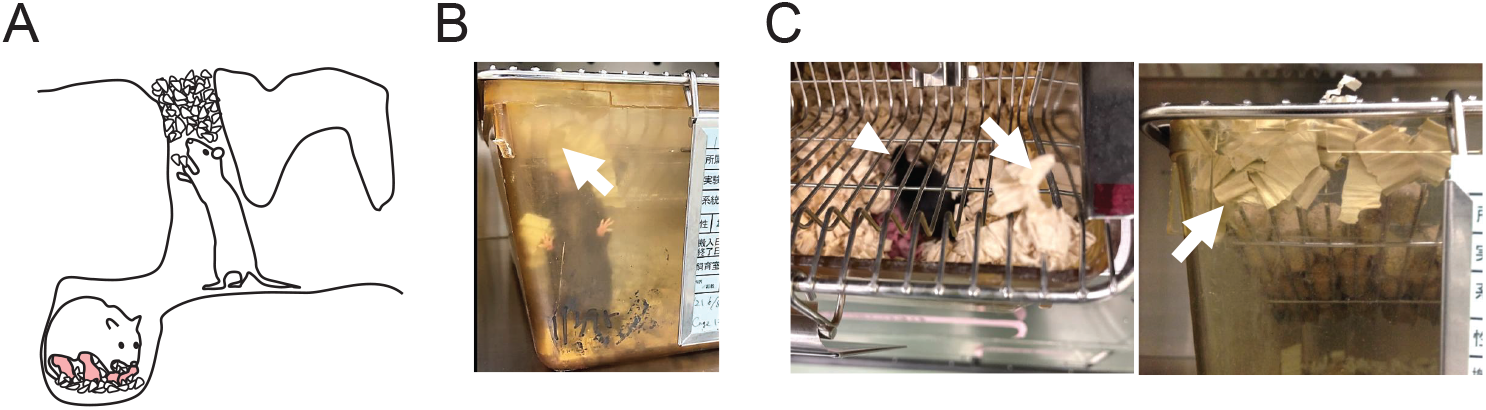
(A) Schematic illustration of entrance-sealing behavior originally described in wild rats (Calhoun, 1963). Lactating females plug the entrance of their burrow with mud, soil, or vegetation to block access by potential intruders while pups remain inside the nest. (B) Example of entrance-sealing behavior induced in laboratory mice by chemogenetic activation of PeFA UCN3 neurons, showing dense packing of bedding material into the cage corner or an entrance-like gap (arrow). (C) Natural entrance-sealing behavior observed in postpartum female mice (arrowhead), in which bedding is actively packed toward small openings or gaps of the home cage (arrows).

Standard behavioral assays therefore provide limited opportunities to examine defensive behaviors that are specifically oriented toward offspring protection. Although sealing-like behavior can be experimentally induced, there has been no reliable method to quantify sealing under controlled laboratory conditions, largely because spontaneous sealing in mice is infrequent and highly variable. Consequently, it has remained difficult to assess how internal state or external threat modulates this form of offspring-centered defense. To address this gap, we developed a simple assay that enables repeated, quantitative measurement of sealing behavior in laboratory mice.

Here, we introduce the Tube-Sealing Test (TST), a simple assay designed to quantify sealing behavior in laboratory mice under standardized conditions. The TST enables repeated and non-invasive measurement of bedding packed into tube-like openings attached to the home cage, providing a reliable readout of sealing intensity. Using this approach, we examined how sex and reproductive state shape spontaneous sealing behavior, and whether sealing in postpartum females is modulated by exposure to an intruder. Together, these experiments establish the TST as a practical tool for investigating offspring-centered defensive behavior in mice. .

## 2. Materials and Methods

### 2-1. Animals

Adult C57BL/6J mice (8–20 weeks old) of both sexes were used. Female mice were tested either as virgins or as postpartum mothers, with postpartum females defined as postpartum day 1–7. Litter size was not experimentally standardized; postpartum females naturally had litters of 5–8 pups. Litters from females showing infanticide or abnormal maternal care were excluded. Prior to behavioral testing, all mice were group-housed. For TST sessions, mice were transferred to individual cages to allow unambiguous quantification of sealing behavior.

All procedures were approved by the Institutional Animal Care and Use Committee of Nara Medical University and were conducted in accordance with national guidelines for the care and use of laboratory animals.

### 2-2. Housing and General Procedures

Mice were maintained in a temperature- and humidity-controlled facility (25°C; 40–60% humidity) under a 12:12 h light/dark cycle (lights on at 08:00). Food and water were available ad libitum. Standard paper-based bedding was used.

For the TST, all mice—males, virgin females, and postpartum females—were transferred to the testing cage 24 h before the first measurement. Postpartum females were moved together with their litter, and pups stayed in the cage continuously. For experiments involving repeated daily measurements, mice remained in the same testing cage across sessions without being returned to group housing.

### 2-3. TST Apparatus

The TST apparatus consisted of a standard mouse home cage fitted with two modified 50-mL plastic centrifuge tubes serving as access tubes. Each tube was prepared by cutting off the conical bottom of a 50-mL tube and wrapping the open end with silicone laboratory tape to create a smooth, safe edge. A circular hole was drilled on each side of the cage wall (11 cm apart), and the modified tubes were inserted through these holes such that the capped ends remained outside the cage while the open ends faced the inside of the cage.

The tubes were secured simply by friction between the taped tube surface and the cage wall, and no additional fasteners were required.

The inner opening of each tube served as the site where bedding could be packed by the mouse. All sealing measurements were performed using the same pair of tubes across sessions.

### 2-4. TST Procedure

At the beginning of the light phase, each TST session was initiated by providing a standardized amount of bedding in the testing cage equipped with the tube apparatus. Mice remained undisturbed for 24 h, after which sealing behavior was quantified. For experiments conducted over multiple consecutive days, this 24 h cycle was repeated without moving mice to a different cage.

### 2-5. Repeated TST measurements for intruder experiments

For intruder experiments, postpartum females underwent three consecutive 24-h TST sessions. On day 0, mice were transferred to the testing cage. Sealing behavior was quantified after each 24-h period on day 1, day 2, and day 3. The two baseline measurements (day 1 and day 2) were used to ensure that sealing behavior was stable before the intruder manipulation. For assessing intruder-induced changes, day 2 was defined as the baseline, and day 3 represented the post-intruder condition.

### 2-6. Scoring of Sealing Behavior

Sealing behavior was quantified by measuring the amount of bedding packed inside the openings of the two cage-attached tubes. After each 24-h TST session, the outer caps of the tubes were unscrewed, and bedding located within the inner opening of each tube was carefully removed using a small curved metal spatula commonly used for weighing milligram-scale powders. The collected bedding was transferred into pre-weighed containers and weighed using an electronic balance. Each tube was measured separately, and the two values were summed to obtain the sealing intensity for each mouse. Only bedding that was clearly located inside the tube opening was considered packed material. Bedding outside the opening or bedding not entering the inner opening was not included. For multi-day experiments, all packed bedding was completely removed after each measurement to ensure that each 24-h session began with an empty tube. The sole quantitative measure used in all analyses was the total packed mass (g).

### 2-7. Statistical Analysis

All statistical analyses were performed using GraphPad Prism 9 (GraphPad Software, CA, USA). Normality of distributions was assessed using the Shapiro–Wilk test. Because sealing intensity data deviated from normality, non-parametric tests were used.

For comparisons of sealing intensity among males, virgin females, and postpartum females, the Kruskal–Wallis’s test was applied, followed by Dunn’s post hoc test for pairwise group comparisons.The occurrence of sealing behavior (sealing vs. no sealing) was analyzed using chi-square tests. When overall group differences were significant, post hoc pairwise comparisons between groups were performed with appropriate adjustment for multiple testing.

For intruder experiments in postpartum females, sealing intensity on the baseline day and on the day following intruder exposure (same individuals) was compared using the Wilcoxon signed-rank test. All tests were two-tailed, and statistical significance was set at p < 0.05.

## 3 Results

### 3-1. Development of the TST

To quantify sealing behavior in laboratory mice, we used the TST, which allows reliable quantification of bedding packed into tube-like openings attached to the home cage (Fig. 2A). The apparatus consists of two modified 50-mL plastic tubes inserted through circular holes in the cage wall, enabling mice to access and manipulate the bedding inside the tube openings. Because the capped ends of the tubes remain outside the cage, packed bedding can be collected and weighed without disturbing the animals. When mice were housed individually in the TST cage for 24-h sessions, both the probability and amount of bedding inserted into the tubes could be measured consistently across sessions (Fig. 2B).

**Figure 2.**
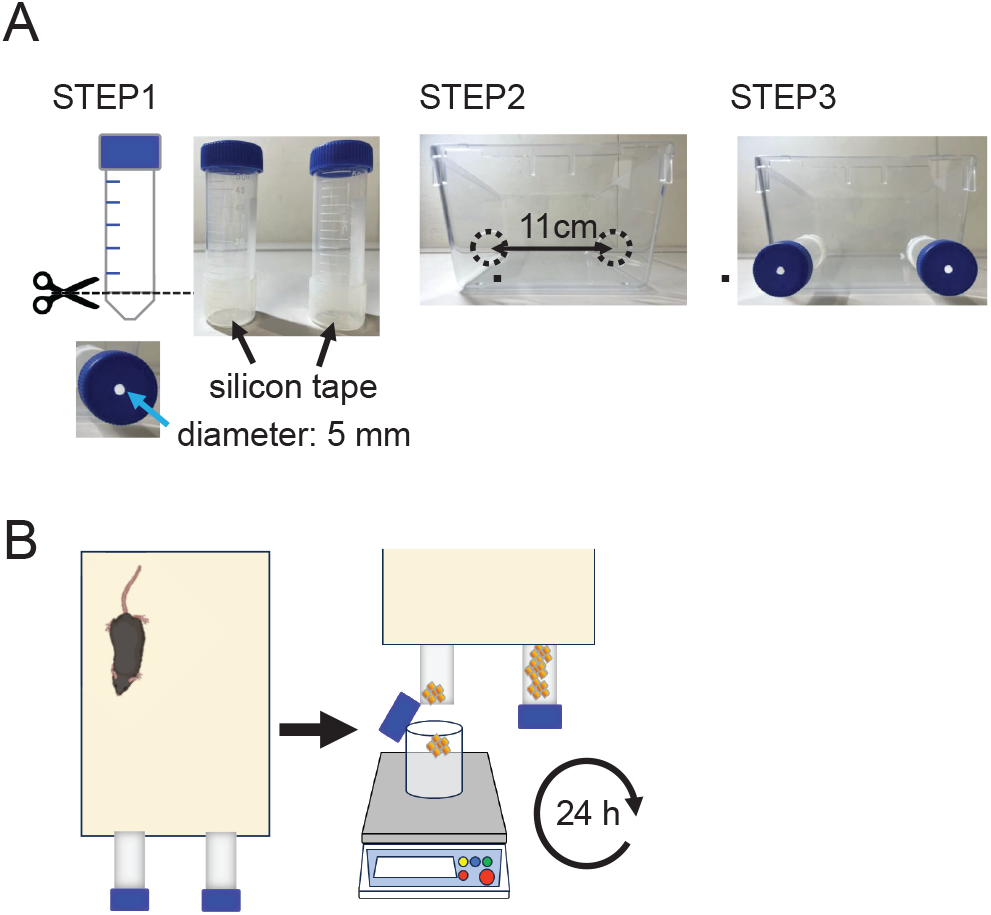
Apparatus and procedure for the TST. (A) Construction of the TST apparatus: STEP 1: The conical bottom of a 50-mL plastic centrifuge tube was removed, and the cut edge was wrapped with silicone tape. A 5-mm-diameter hole was made in the tube cap. STEP 2: Two circular holes (11 cm apart) were drilled in the side wall of a standard mouse cage. STEP 3: The modified tubes were inserted through the holes, with the capped ends positioned outside the cage. (B) Procedure for quantifying sealing behavior. Mice were housed individually in the TST cage for 24 h. After the session, the outer caps were unscrewed and bedding packed inside the tube openings was collected and weighed to determine the sealing intensity.

### 3-2. Sex and reproductive state differences in tube-sealing behavior

Representative images of tube-sealing behavior illustrated that males rarely inserted bedding into the tubes, whereas virgin females occasionally packed bedding, and postpartum females displayed pronounced tube-sealing behavior (Fig. 3A). Heatmaps showing sealing intensity across repeated daily sessions for each individual revealed clear group differences (Fig. 3B). Males showed little to no sealing across sessions. Virgin females exhibited moderate sealing with substantial individual variability, whereas postpartum females consistently showed strong sealing behavior.

**Figure 3.**
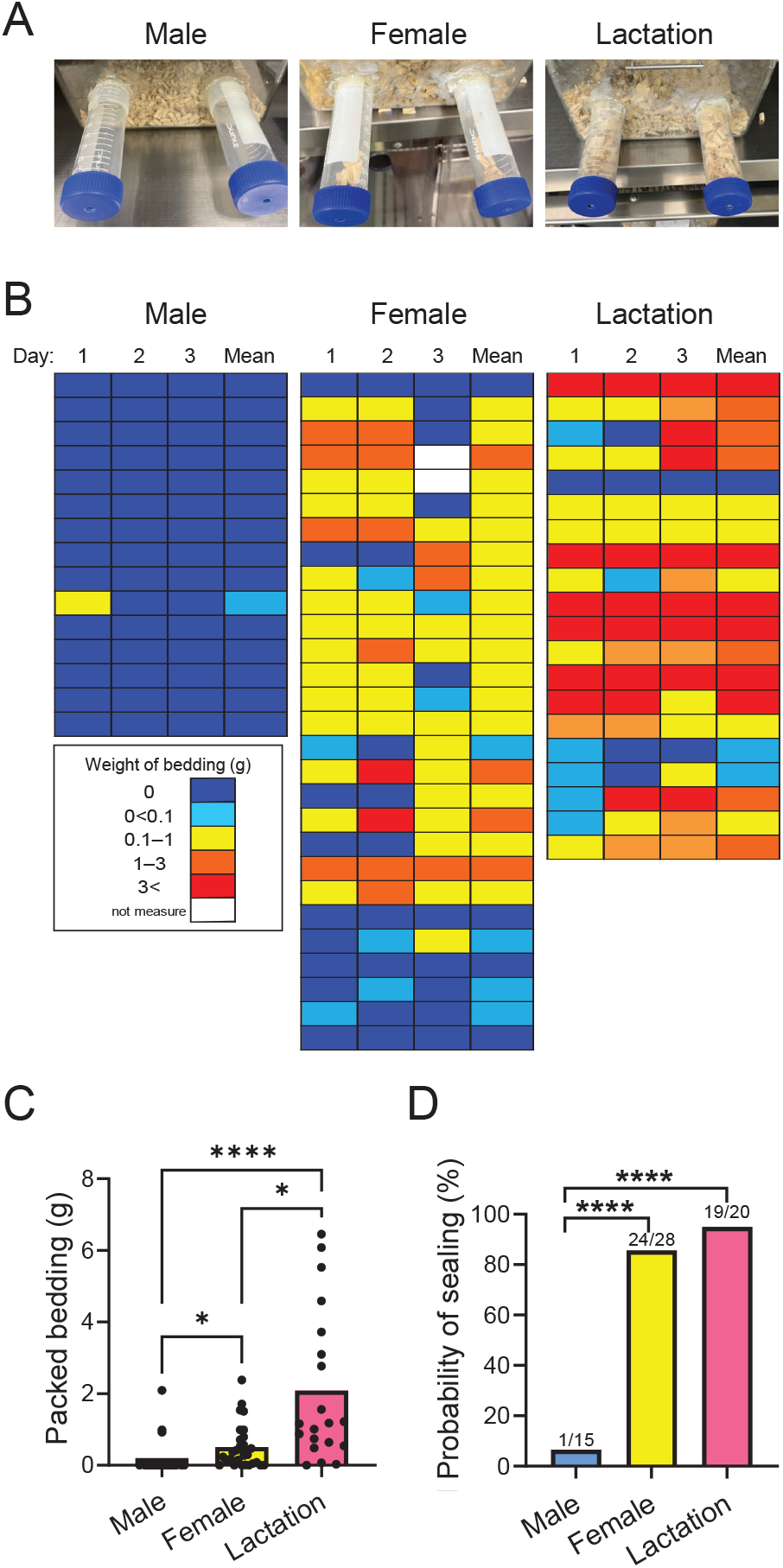
Sex and reproductive state differences in tube-sealing behavior. (A) Representative examples of tube-sealing behavior in males, virgin females, and postpartum (lactating) females. Males rarely packed bedding into the tubes, whereas females—especially postpartum females—frequently inserted bedding material. (B) Heatmaps showing tube-sealing intensity across repeated daily sessions for each individual. Blue indicates no sealing (0 g), and warmer colors (yellow → orange → red) represent greater amounts of packed bedding. Males showed almost no sealing across sessions. Virgin females showed moderate sealing with large individual variability. Postpartum females exhibited robust tube-sealing behavior. (C) Probability of tube-sealing (percentage of animals that sealed at least once across the measured sessions). Postpartum females (19/20) and virgin females (24/28) showed a significantly higher probability of sealing than males (1/15) (χ^2^ test, ****p < 0.0001). (D) Amount of packed bedding (g) averaged across the measured sessions. Postpartum females packed significantly more bedding than virgin females and males (Kruskal–Wallis test with Dunn’s post hoc test; *p < 0.05, ****p < 0.0001).

The probability of tube-sealing—defined as sealing at least once during the measured sessions—also differed across groups. Only 1 of 15 males sealed at any point, compared with 24 of 28 virgin females and 19 of 20 postpartum females (χ^2^ test, ****p < 0.0001) (Fig. 3C). The amount of bedding packed into the tubes likewise varied across groups. Postpartum females inserted significantly greater amounts of bedding than virgin females and males (Kruskal–Wallis test with Dunn’s post hoc test; *p < 0.05, ****p < 0.0001) (Fig. 3D).

### 3-3. Intruder exposure selectively enhanced tube-sealing behavior in postpartum females

To test whether sealing behavior is modulated by a direct threat experience, we examined the effects of intruder exposure in postpartum females (Fig. 4). Following two baseline sessions (Day 1–2), a male intruder was introduced through one of the tubes on Day 2, and sealing behavior was quantified again on Day 3. Sealing intensity on the tube not exposed to the intruder showed no significant change between pre-intruder and post-intruder sessions (Fig. 4B, Intruder −). In contrast, sealing intensity on the intruder-exposed tube increased significantly after intruder exposure (Wilcoxon signed-rank test, ***p < 0.001) (Fig. 4B, Intruder +). Total sealing across both tubes was also significantly elevated following intruder exposure (**p < 0.01). These results demonstrated that tube-sealing behavior in postpartum females was modulated by direct exposure to an intruder.

**Figure 4.**
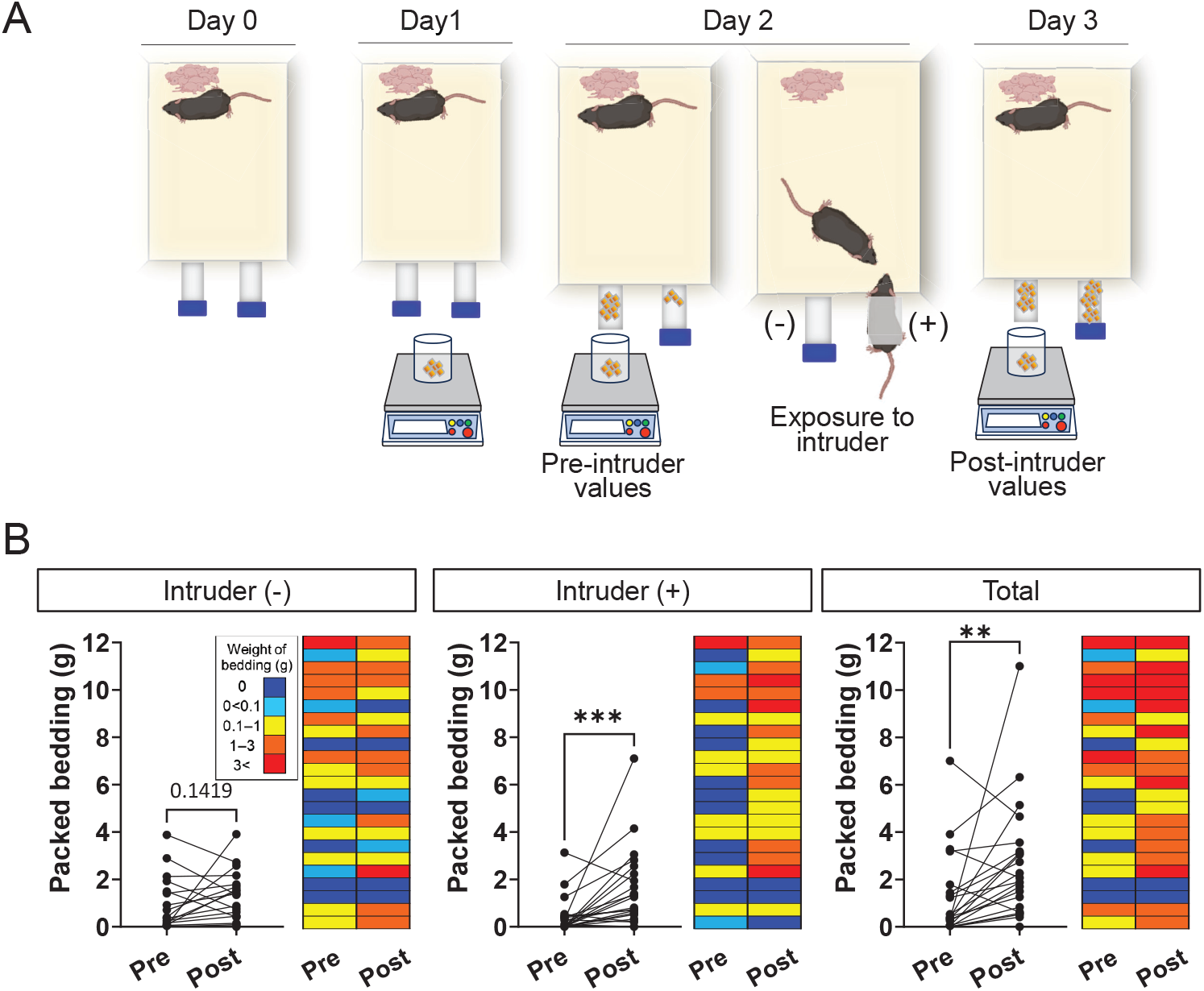
Intruder exposure enhances tube-sealing behavior in postpartum females. (A) Experimental timeline for the intruder assay. Postpartum females were transferred to the TST cage on Day 0. Tube-sealing behavior was quantified on Day 1 and Day 2 to obtain pre-intruder values. On Day 2, a male intruder was introduced through one of the tubes, allowing brief direct contact before release into the cage. Sealing behavior was quantified again on Day 3 to obtain post-intruder values. (B) Effects of intruder exposure on tube-sealing behavior. **Left:** Tubes on the side opposite the intruder (Intruder −) showed no significant change in sealing intensity between pre- and post-intruder sessions (Wilcoxon signed-rank test; ns, p = 0.1419). **Middle:** Tubes on the intruder-exposed side (Intruder +) showed a significant increase in sealing intensity following intruder exposure (***p < 0.001). **Right:** Total sealing across both tubes significantly increased after intruder exposure (**p < 0.01). Heatmaps adjacent to each graph illustrate sealing intensity for each individual mouse before (Pre) and after (Post) intruder exposure. Blue indicates no sealing (0 g), whereas progressively warmer colors (yellow → orange → red) represent greater amounts of packed bedding.

## 4. Discussion

In this study, we established the TST as a simple assay that enables reliable quantification of bedding-packing behavior in mice. Using this assay, we demonstrated that tube-sealing behavior was strongly sex-biased, being virtually absent in males and expressed most robustly in postpartum females. Although virgin females occasionally engaged in sealing, postpartum females showed markedly greater sealing intensity. Furthermore, exposure to an intruder selectively enhanced tube-sealing behavior in postpartum females, indicating that this behavior is sensitive to direct threat experiences. Together, these findings identify tube-sealing as a reproducible and ethologically relevant defensive behavior that is amplified during the postpartum period.

Tube-sealing observed in the TST shares key features with entrance-sealing behaviors described in wild rodents(Calhoun, 1963), in which lactating females block access points to protect pups from potential intruders. Similar sealing-like behaviors have also been observed in laboratory mice when PeFA UCN3 neurons are experimentally activated, suggesting that the motor pattern for sealing is already embedded within the neural repertoire of the species (Horii-Hayashi et al., 2023). The present results extend these observations by demonstrating that postpartum females spontaneously express sealing behavior in a controlled laboratory environment. Thus, tube-sealing captured by the TST represents an ethologically relevant defensive action that can be quantified under standardized conditions.

Although both virgin and postpartum females frequently engaged in tube-sealing, postpartum females displayed a markedly greater sealing intensity. The lack of a statistical difference in the occurrence of sealing between virgin and postpartum females indicates that the propensity to initiate sealing is already present in non-maternal females. In contrast, the magnitude of sealing was strongly enhanced during the postpartum period. This dissociation suggests that reproductive state does not alter the likelihood of producing sealing behavior, but selectively increases the vigor or amount of sealing once the behavior is initiated. Such state-dependent amplification is consistent with postpartum behavioral adaptations that strengthen defensive responses without modifying their basic probability of occurrence.

Although some postpartum females displayed aggressive behaviors toward the intruder—such as chasing, anogenital investigation, or occasional attacks—these responses are consistent with forms of maternal aggression commonly observed in lactating rodents (Lonstein and Gammie, 2002). However, because these behaviors were not quantified in a standardized manner, we refrain from drawing conclusions about their relationship to sealing. Importantly, the increase in tube-sealing did not appear to co-occur consistently with clear aggressive acts, suggesting that tube-sealing reflects a distinct defensive mode rather than a byproduct of heightened arousal. Whereas maternal aggression represents a reactive defense that is expressed once a threat is already present, tube-sealing embodies a preventive and anticipatory form of offspring-centered protection. By creating a physical barrier before direct contact occurs, sealing behavior functions as a proactive strategy that limits potential access to offspring. Together, these observations indicate that tube-sealing constitutes a proactive, predictive mode of maternal defense that is functionally distinct from the reactive aggression typically expressed during postpartum threat encounters.

The tube-selective increase in sealing after intruder exposure suggests that sealing intensity may reflect a form of context-dependent defensive motivation. Although the present study does not identify the underlying mechanisms, the specificity of the response raises the possibility that postpartum females modulate sealing intensity through neural or endocrine pathways that enhance defensive responding when offspring may be at risk. This assay therefore offers a means to examine how threat-related cues shape maternal defensive strategies.

A methodological limitation of the TST is that its applicability is constrained to the early postpartum period. As pups become increasingly mobile, they occasionally enter the tubes and disturb the bedding, making it difficult to determine whether packed material reflects maternal behavior or pup activity. For this reason, sealing measurements obtained beyond the early postpartum phase may not accurately represent maternal sealing intensity. Future refinements of the assay will be necessary to extend its use to later stages of the postpartum period or to conditions in which pup interference can be minimized.

In summary, the TST provides a simple approach for quantifying tube-sealing behavior and reveals that sealing intensity is strongly enhanced during the postpartum period and further amplified by exposure to an intruder. These findings identify tube-sealing as a measurable component of offspring-centered defense that can be assessed under standardized laboratory conditions. By enabling repeated, non-invasive measurements, the TST may facilitate future studies aimed at uncovering how internal state and external threats interact to shape maternal defensive behavior.

## 6. Author Contributions

NH-H designed the study, performed experiments, analyzed data, and wrote the manuscript. SM, HT, and KA assisted with experiments and data analysis. KI supervised the project and contributed to manuscript editing. All authors approved the final version of the manuscript.

## 8. Acknowledgements

We would like to thank Michiko Kitsuki and Yumi Moriwake for their support with our experiments. We also thank the animal facility staff for excellent animal care. Some illustrations were created with BioRender.com.

## 9. Data Availability

The datasets generated for this study are available from the corresponding author upon reasonable request.

